# ZEB1 regulates BCL2 in cancer-associated fibroblasts to promote cholangiocarcinoma chemoresistance to gemcitabine and cisplatin

**DOI:** 10.1101/2025.09.10.675294

**Authors:** Ester Gonzalez-Sanchez, Marie Vallette, Aashreya Ravichandra, Mirko Minini, Allan Pavy, Josep Amengual, Carlos Andres Roldán-Hernández, Corentin Louis, Juan Jose Lozano, Isabel Fabregat, Nathalie Guedj, Valérie Paradis, Cedric Coulouarn, Lynda Aoudjehane, Laura Fouassier, Javier Vaquero

## Abstract

Intrahepatic cholangiocarcinoma (iCCA) is an agressive tumour from the biliary tree that is characterized by a prominent desmoplastic stroma mainly composed of cancer-associated fibroblasts (CAF) and a poor prognosis due to its late clinical presentation and the lack of effective non-surgical treatments. Current therapies still include chemotherapeutic combinations of gemcitabine and cisplatin for the majority of the patients showing poor results due the apparition of resistance. This situation led us to interrogate the potential role in the development of chemoresistance of ZEB1, an EMT-inducing transcription factor (EMT-TF) that we previously identified as a pro tumorigenic factor in tumour cells and CAF of iCCA. By analysing human CCA samples and sc-RNAseq public databases, we show here that ZEB1 is present in the tumour microenvironment of all iCCA patients tested and is prominently expressed by CAF. Using cellular models of CAF, we show that cells depleted for ZEB1 are more sensitive to gemcitabine and cisplatin, via a mechanism involving the regulation of the anti-apoptotic gene BCL2. Moreover, ZEB1 expressing CAF protect iCCA tumour cells against the toxicity of the chemotherapeutic drugs, an event that could be reversed by a BCL2 inhibitor venetoclax. Therefore, our results point to the use of BCL2 inhibitors to improve the efficacy of current chemotherapeutic regimens of gemcitabine and cisplatin in iCCA patients.

## 1. Introduction

Intrahepatic cholangiocarcinoma (iCCA) is a very aggressive malignancy from the biliary tree and represents the second most common primary liver malignancy after hepatocellular carcinoma (HCC).[1, 2] Late diagnosis often compromises surgical resection, which is the only effective therapeutic option but is applicable in only 20% of cases.[1, 2] Moreover, the lack of specific etiologic factor in most of the cases impedes the surveillance of a population with predisposing conditions, contributing to the poor prognosis of the disease.[1, 2] For the majority of iCCA patients who are not eligible to undergo surgery or who do not meet the criteria for targeted therapies, the combination of chemotherapy (gemcitabine plus cisplatin) with immunotherapy (durvalumab or pembrolizumab) was recently approved as the new standard of care.[3, 4] Nevertheless, the absence of factors to better stratify the patients and the intrinsic chemoresistance of iCCA contributes to the lack of efficacy of this chemotherapeutic combination in an important number of patients. Therefore, it is essential to investigate and characterize new actionable targets for the treatment of this deadly disease.

iCCA is characterized by a prominent tumour microenvironment (TME) enriched in cancer associated fibroblasts (CAF) that produce abundant collagen and elastic fibres deposited in the extracellular matrix (ECM), contributing to tumour progression and chemoresistance.[1, 2] In recent years, the TME, and more specifically CAF have received a significant attention as potential therapeutic targets to combat iCCA progression.[5] Indeed, our previous study showed that CAF express ZEB1, an EMT-inducing transcription factor (EMT-TF), which regulates the activation status of CAF and their interplay with tumour cells through the release of soluble factors.[6] In many tumours, ZEB1 expression in cancer cells has been linked to resistance to a plethora of chemotherapeutic drugs, including gemcitabine and cisplatin.[7-9] However, ZEB1 role in the TME-induced chemoresistance is still largely unknown.

Here we aimed to investigate the impact of ZEB1 in the tumour resistance induced by CAF to current chemotherapeutic regimens in iCCA. By analysing sc-RNAseq data and iCCA patient samples, we demonstrate that ZEB1 is always present in CAF from the TME. Furthermore, our thorough *in vitro* experimentation reveals that: 1/ ZEB1 regulates BCL2 inducing a resistance to gemcitabine and cisplatin in 2D cellular models of CAF; 2/ CAF reduce the sensibility of malignant cells to chemotherapeutic drugs in a 3D mixed tumour cells-CAF spheroid model. Interestingly, the BCL2 inhibitor venetoclax restores the sensitivity to gemcitabine and cisplatin.

## 2. Material and methods

### 2.1. Reagents

Gemcitabine (G6423, Sigma-Aldrich), cisplatin (479306, Sigma-Aldrich) and sorafenib (BAY 43-9006, Sellechchem) were used at the concentrations indicated in the figures and figure legends. TGF-β1 (#T70039, Sigma-Aldrich) was used at 2 ng/mL. Venetoclax (HY-15531, MedChemExpress, Monmouth Junction, NJ, USA) was used at 5 μM. Puromycin (A11138-03, Life Technologies, Carlsbad, CA, USA) was used at 4 µg/mL. Hygromycin (ant-hg-1, InvivoGen, San Diego, CA, USA) was used at 100 µg/mL.

### 2.2. Cell culture and treatment

The human hepatic stellate cell (HSC) lines hTERT-HSC and LX2-HSC were kindly provided by Dr. L. Aoudjehane (ICAN, Paris, France). hTERT-HSC and LX2-HSC were cultured in DMEM 4.5 g/L glucose (31966-021). HuCCT1 cells, derived from iCCA, were kindly provided by Dr. G. Gores (Mayo Clinic, MN). SK-ChA-1 cells, derived from extrahepatic CCA, were obtained from Dr. A. Knuth (Zurich University, Switzerland). HuCCT1 and SK-ChA-1 cells were cultured in DMEM 1 g/L glucose (21885-025). All media were supplemented with antibiotics (100 UI/mL penicillin and 100 μg/mL streptomycin (15140-122), antimycotic (0.25 mg/mL amphotericin B (15290018), and 10% FBS (10270-106). Cell lines were routinely screened for the presence of mycoplasma and authenticated for polymorphic markers to prevent cross-contamination.

### 2.3. Gain and loss-of-function cellular models

#### Stable ZEB1 inhibition

Stable ZEB1 inhibition in SK-ChA-1 and liver myofibroblasts (hTERT-HSC cells) was previously performed by lentiviral transduction with specific small hairpin RNAs (shRNAs).[6] Plasmids containing sequences targeting ZEB1 (shRNA-ZEB1) or scramble (shRNA-Control), designed by Genecopoeia, were used to produce ΔU3 SIN lentivirus in the Viral and Gene Transfer Vectors Platform of Necker (Paris, France). Here we achieved stable ZEB1 inhibition in LX2-HSC cells following the same method. Briefly, cells were infected and maintained in the presence of lentivirus for 24 hours. Then, lentiviruses were removed, and fresh media was added to maintain cells in culture another 48 hours until mCherry expression was observed. Positive transduced cells were detected by positive fluorescence signal, and they were selected by treatment with lethal doses of puromycin for 72 hours to eliminate noninfected cells. Due to incomplete efficiency of puromycin selection in SK-ChA-1 cells, two clones of each cell type were isolated and characterized in our previous study.[6]

#### Stable BCL2 overexpression

Stable BCL2 overexpression in hTERT-HSC was performed by lentiviral transduction. Lentivirus encoding BCL2 (NM_000633.3) were obtained from Genecopoeia. We used the same protocol than above, but positive cells were identified by eGFP expression and were selected with lethal doses of hygromycin.

### 2.4. Gene expression profiling

*TCGA cohort:* The mRNA expression levels of *ZEB1* versus *BCL2* and *BAX* in CCA tumours from the repository of The Cancer Genome Atlas (TCGA) were analysed through the cBioPortal database (https://www.cbioportal.org/) and represented as Spearman correlation coefficients (r) and P-values to measure the significance.[10, 11]

*Rennes cohort*: gene expression datasets established from whole[12] or laser microdissected[13] iCCA tumours were previously described. Clinical and pathological features of a cohort of 39 patients with iCCA (thereafter referred to as the Rennes cohort) were previously reported.[13] Freshly frozen human tumour samples were obtained from the processing of biological samples through the Centre de Ressources Biologiques (CRB) Santé of Rennes (BB-0033-00056) and the French liver cancer biobanks network – INCa (BB-0033-00085). The research protocol was conducted under French legal guidelines and fulfilled the requirements of the local institutional ethics committee.

*single-cell RNA-seq*: the single-cell RNA-seq dataset was processed, analysed and visualized using the Trailmaker® (https://scp.biomage.net/) hosted by Biomage (https://biomage.net/). First, pre-filtered count matrices from the Shi et. al [GSE201425] were uploaded to Trailmaker®. Briefly, to pre-process the data, barcodes with low unique molecular identifiers (UMIs) and or high mitochondrial reads were filtered out. Next, to exclude outliers, a robust linear model was fitted to all samples. Lastly, barcodes with high doublet scores were excluded from the analysis, resulting in high quality barcodes that was used in subsequent integration steps. Briefly, to integrate the barcodes, data was log-normalized and the top 2000 highly variable genes (HSVs) were selected. Next, principal component analysis (PCA) was performed, and the top 30 principal components, explaining 89.67% of the total variance was used to perform sample batch correction with the integrated Harmony R package. Louvain method of clustering was employed. To reduce dimensionality, data was visualized using t-distributed stochastic neighbour embedding (t-SNE). To identify cluster-specific marker genes, marker genes of each cluster was compared to all other clusters and appropriately annotated.

### 2.5. RNA and reverse transcription-PCR

Total RNA extraction and RT-qPCR were performed as previously described.[14] Primer sequences are provided in Supplementary Table 1. Gene expression was normalized to *GAPDH* mRNA and was expressed relatively to the control condition of each experiment. The relative expression of each target gene was determined from replicate samples using the formula 2^-ΔΔCt^.

### 2.6. Chromatin immunoprecipitation PCR (ChIP-PCR) analysis

We uploaded the supplementary files (bigWig and narrowPeaks) involved in Durant et al publication[15] from GEO[16] (https://www.ncbi.nlm.nih.gov/geo/query/acc.cgi?acc=GSE246672) and we show where ZEB1 binding peaks at BCL2 gene (genome boundaries as chr18:63123346-63320769) using UCSC genome browser[17] (viewLimitsMax 0:0.5).

### 2.7. Immunoblot analysis

For obtaining whole-cell lysates for immunoblotting, cell cultures were lysed in RIPA buffer (Sigma) supplemented with 1 mmol/L orthovanadate and a cocktail of protease inhibitors. Proteins were quantified using a BCA kit (Pierce). Western blot analysis was performed as previously described.[14] Primary antibodies are provided in Supplementary Table 2.

### 2.8. Immunofluorescence

Immunofluorescence assays were performed as previously described.[14] The list of primary antibodies is provided in Supplementary Table 4. Cells were observed with an Olympus Bx 61 microscope (Olympus).

### 2.9. Viability assays

Cells were plated in 96-well plates. Twenty-four hours later, the medium was replaced by fresh culture medium in the absence (vehicle) or presence of the corresponding drugs, as indicated in the figures and figure legends. Cells were then incubated for 72 h before determining the viability by the crystal violet method. Absorbance was quantified with a spectrophotometer (Tecan) at 595 nm.

### 2.10. Sphere formation assay

Spheroids were generated by the hanging drop method. For all the experiments, cells were suspended, at a concentration of 300.000 cells/mL for HuCCT1 tumour cells alone or 300.000 + 300.000 for mixed spheroids containing HuCCT1 cells and hTERT-HSC cells in a proportion 1:1. Cells were suspended in medium with 0.24% of methylcellulose (M7027, Sigma-Aldrich) in the absence or presence of the corresponding drugs at the concentrations indicated in the figures and figure legends. Twenty-five µl drops were pipetted onto the lid of 100 mm dishes, that were inverted over dishes containing 5 ml of cell culture medium to avoid drying. After 4 days of incubation at 37°C and 10% CO_2_, spheroids were transferred by pipetting onto a low-attachment 6-well culture plate (3471, Corning) and incubated another 4 days with fresh medium and drugs. Then, images of 10–20 spheroids per experiment were acquired with a NIKON Eclipse Ti2 microscope and spheroids size was measured with ImageJ software (National Institute of Health, USA).

### 2.11. CCA specimens

We retrospectively retrieved from the files of surgical and pathology department patients who had undergone liver resection for iCCA between 2002 and 2014 at Beaujon Hospital, Clichy, France. All clinicopathological and follow up data of patients were collected and registered in data base (Supplementary Table 4). Informed consent was obtained in all cases and the study protocol conformed to the ethical guidelines of the 1975 Declaration of Helsinki as reflected in a priori approval by the appropriate institutional review committee.

### 2.12. Immunohistochemical analysis on Tissue MicroArray (TMA)

Paraffin embedded tissue blocks were used for tissue microarray construction. The slides were reviewed to identify and mark representative areas of viable tumour tissue. Taking tumour heterogeneity into account, three tissue cores of 1mm each were punched from selected from selected tumour areas of any donor tissue block and brought into a recipient paraffin block. We used a tissue arraying instrument (MTA-1, Beecher Instrument, Inc., Sun Prairie, WI, USA). A total of five TMA were built.

Co-immunostaining between ZEB1 and alpha-smooth muscle actin (α-SMA) according the ultraview double immunostained protocol (optiview DAB/ultraview RED) (Supplementary Table 3). All sections were evaluated by pathologists (NG and VP) and a researcher (JV) using a semi-quantitative analysis. ZEB1 was considered positive when nuclear staining in tumour cells was observed whatever the intensity. α-SMA was considered positive when CAF were immunostained.

### 2.13. Statistical analysis

Results were analysed using the GraphPad Prism 5.0 statistical software. Data are shown as means ± standard error of the mean (SEM). For comparisons between two groups, parametric Student t test or nonparametric Mann–Whitney test were used. For comparisons between more than two groups, parametric one-way ANOVA test followed by a posteriori Bonferroni test was used.

## 3. Results

### 3.1. ZEB1 is widely expressed in CAF from iCCA

We previously showed that α-SMA positive CAF from iCCA samples and HSC derived cell lines expressed ZEB1.[6] To widen our study on ZEB1, we carried out a more detailed study by analysing ZEB1 expression in a wider set of human iCCA samples by IHC and by interrogating public datasets of sc-RNAseq. The IHC exploration shows that ZEB1 is expressed by α-SMA positive CAF in 100% of iCCA samples analysed, independently of the tumour differentiation (Figure 1A). sc-RNAseq analysis identified ZEB1 expression in multiple cell subtypes, including tumour cells, immune cells and endothelial cells, but showed that CAF population exhibited the highest levels of ZEB1 expression (Figure 1B). To further analyse the role of ZEB1 *in vitro*, we took advantage of a CAF-like cellular model derived from human HSC, hTERT-HSC, downregulated for ZEB1 that we previously generated by the means of lentiviral infection.[6] Here we used the same procedure to generate another CAF-like cellular model from LX2-HSC in which ZEB1 is downregulated (Supplementary Figure 1A-C). Further comparison with human primary CAF isolated from iCCA samples showed a clear nuclear localization of ZEB1 (Figure 1C). Moreover, α-SMA expression decreased gradually in parallel with that of ZEB1 from iCCA-CAF (highest expression, left panel) to LX2-HSC-shControl (lowest expression, right panel) (Figure 1C). Downregulation of ZEB1 in both CAF models (hTERT-HSC-shZEB1 and LX2-HSC-shZEB1) results in loss of a-SMA staining compared with shControl cells (Figure 1C), validating our previous observations that ZEB1 regulates the basal activation state of liver HSC and CAF.[6] Altogether, our data demonstrates that ZEB1 is widely expressed both in native CAF from iCCA and in CAF-like models derived from HSC, suggesting an important role in the functionality of these cells.

**Figure 1.**
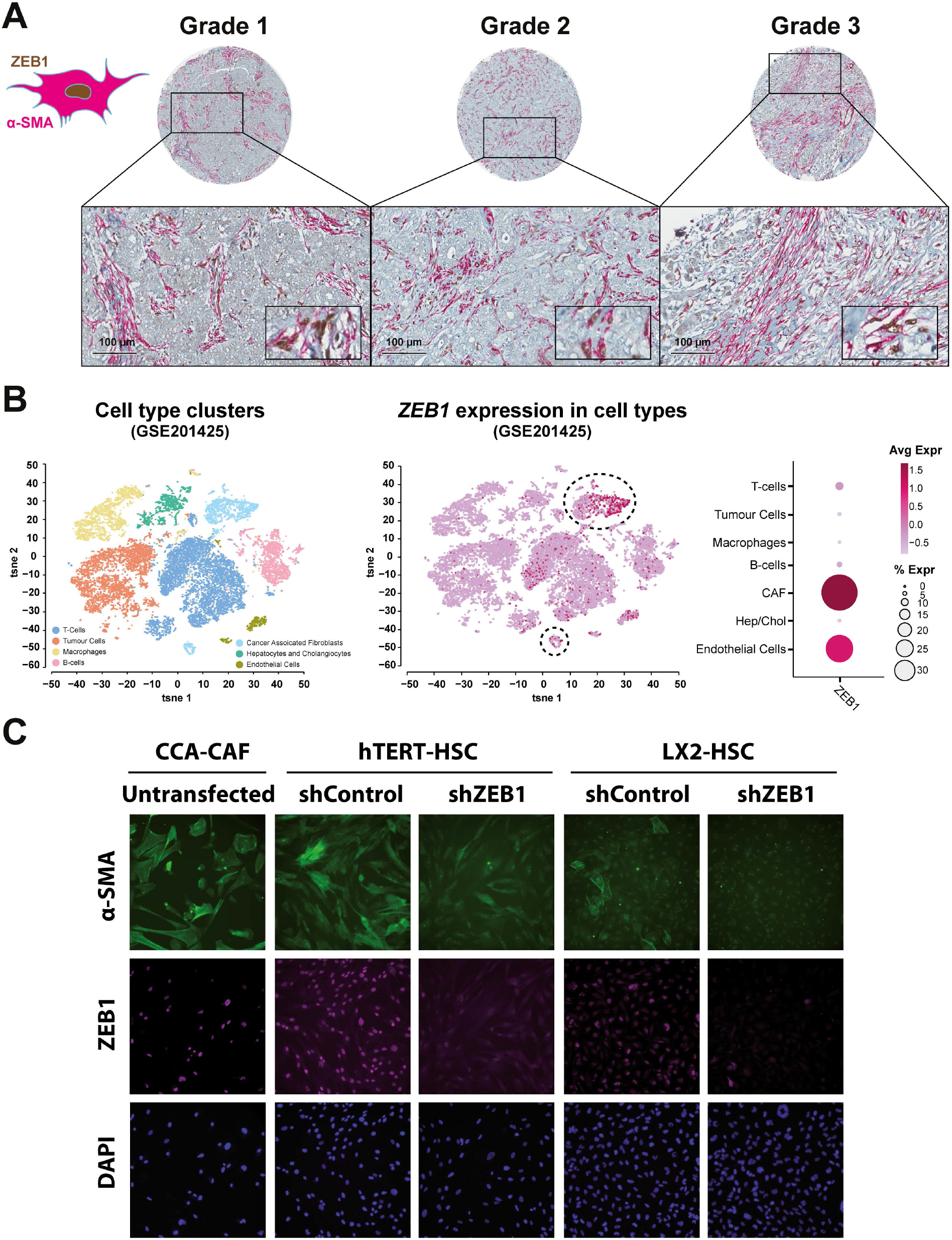
Cancer-associated fibroblasts (CAF) from human intrahepatic cholangiocarcinoma (iCCA) express ZEB1. **A**. Representative IHC co-immunostaining of ZEB1 (brown) and α-SMA (pink) in human iCCA showing ZEB1-expressing CAF. ZEB1 expression was found in CAF in all of the 45 samples analysed. **B**. T-SNE and dotplot showing ZEB1 expression in sc-RNAseq data set GSE201425 from CCA biopsies. **C**. Representative images of α-SMA (green) and ZEB1 (purple) expression in shRNA-Control and shRNA-ZEB1 hTERT-HSC and LX2-HSC cells analysed by immunofluorescence. Nuclei (in blue) were labelled with DAPI. Comparison with CAF isolated from human iCCA (left panel) is shown.

### 3.2. ZEB1 down-regulation sensitizes CAF to gemcitabine and cisplatin

Most iCCA patients still receive traditional chemotherapeutic agents, including gemcitabine and cisplatin.[18] In addition, ZEB1 has been related to increase chemoresistance to these drugs in tumour cells from other cancers.[7-9] Thus, we performed viability studies in both CAF models, hTERT-HSC and LX2-HSC, to test if that was also the case in these cells. Indeed, ZEB1 downregulation significantly increased CAF sensitivity to both gemcitabine and cisplatin (Figure 2A-F). The IC50 of both drugs was reduced in both hTERT-HSC and LX2-HSC cells depleted for ZEB1 (Figure 2C and F), although the reduction of gemcitabine IC50 in LX2-HSC-shZEB1 cells was less pronounced because LX2-HSC cells are already very sensitive to this drug (Figure 1C). Interestingly, the IC50 of cisplatin (Figure 1F) correlated perfectly with the expression of ZEB1 in these cell lines (Supplementary Figure 1B). Surprisingly, ZEB1 downregulation did not alter the sensitivity of CCA tumour cells to either gemcitabine or cisplatin (Supplementary Figure 2). Similarly, we did not observe changes in the response of liver myofibroblasts to sorafenib (Supplementary Figure 3), one of the major drugs given to hepatocellular carcinoma (HCC) patients, where HSC also give rise to CAF during hepatocarcinogenesis. These results indicate that ZEB1 plays a specific role in regulating the resistance of CAF to the chemotherapeutic agents administrated to patients with iCCA.

**Figure 2.**
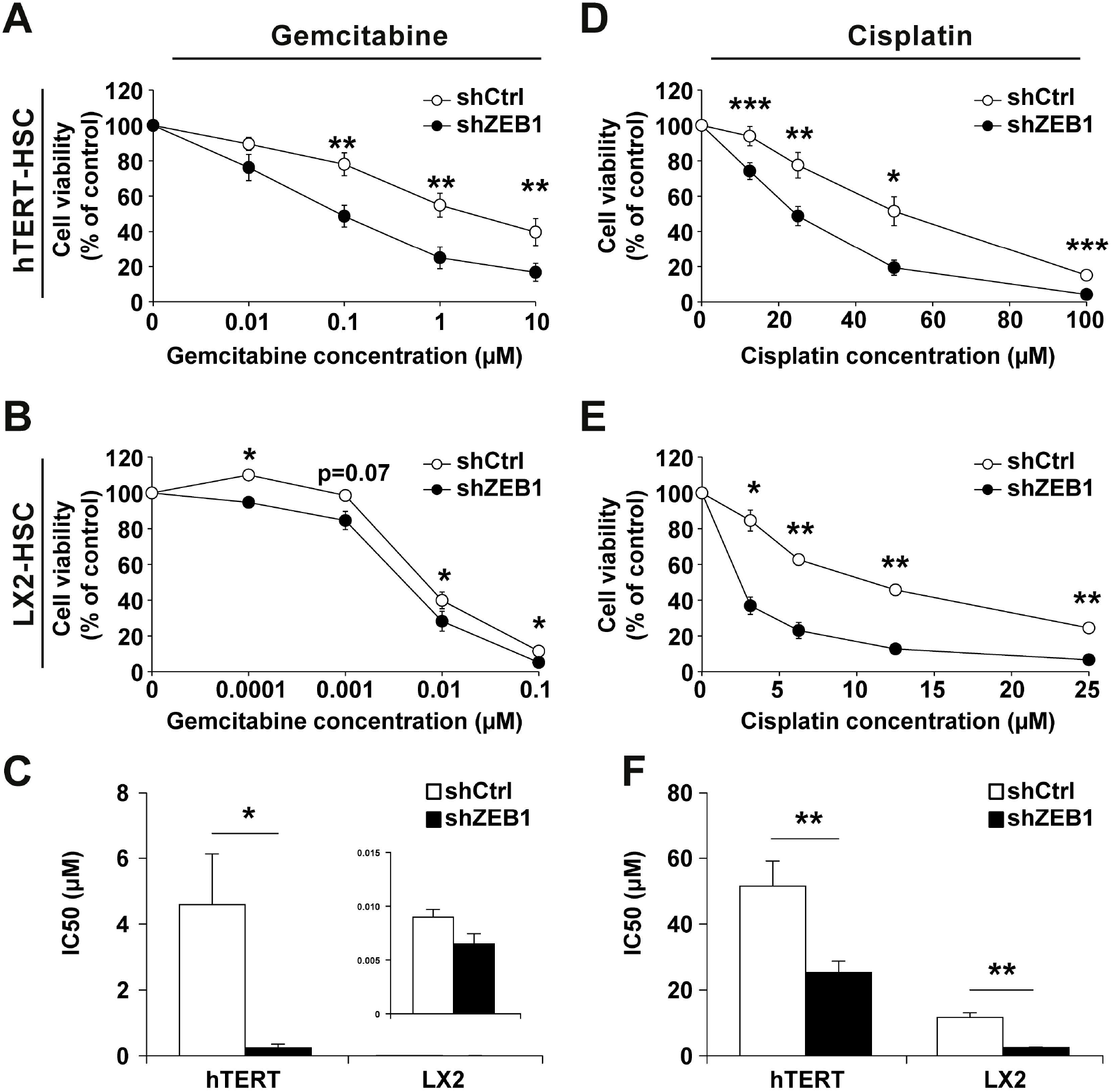
ZEB1 protects liver myofibroblasts against the toxicity of chemotherapeutic drugs used for the treatment of intrahepatic cholangiocarcinoma (iCCA). Effect of gemcitabine (**A-C**) and cisplatin (**D-F**) on the viability of shRNA-Control (shCtrl) and shRNA-ZEB1 hTERT-HSC and LX2-HSC cells. Cell viability was measured by the crystal violet test after incubation with the indicated concentrations of gemcitabine or cisplatin for 72h. Values are expressed as means ± SEM from at least 5 cultures. *, p < 0.05; **, p < 0.01; ***, p < 0.001; as compared with shCtrl cells.

**Figure 3.**
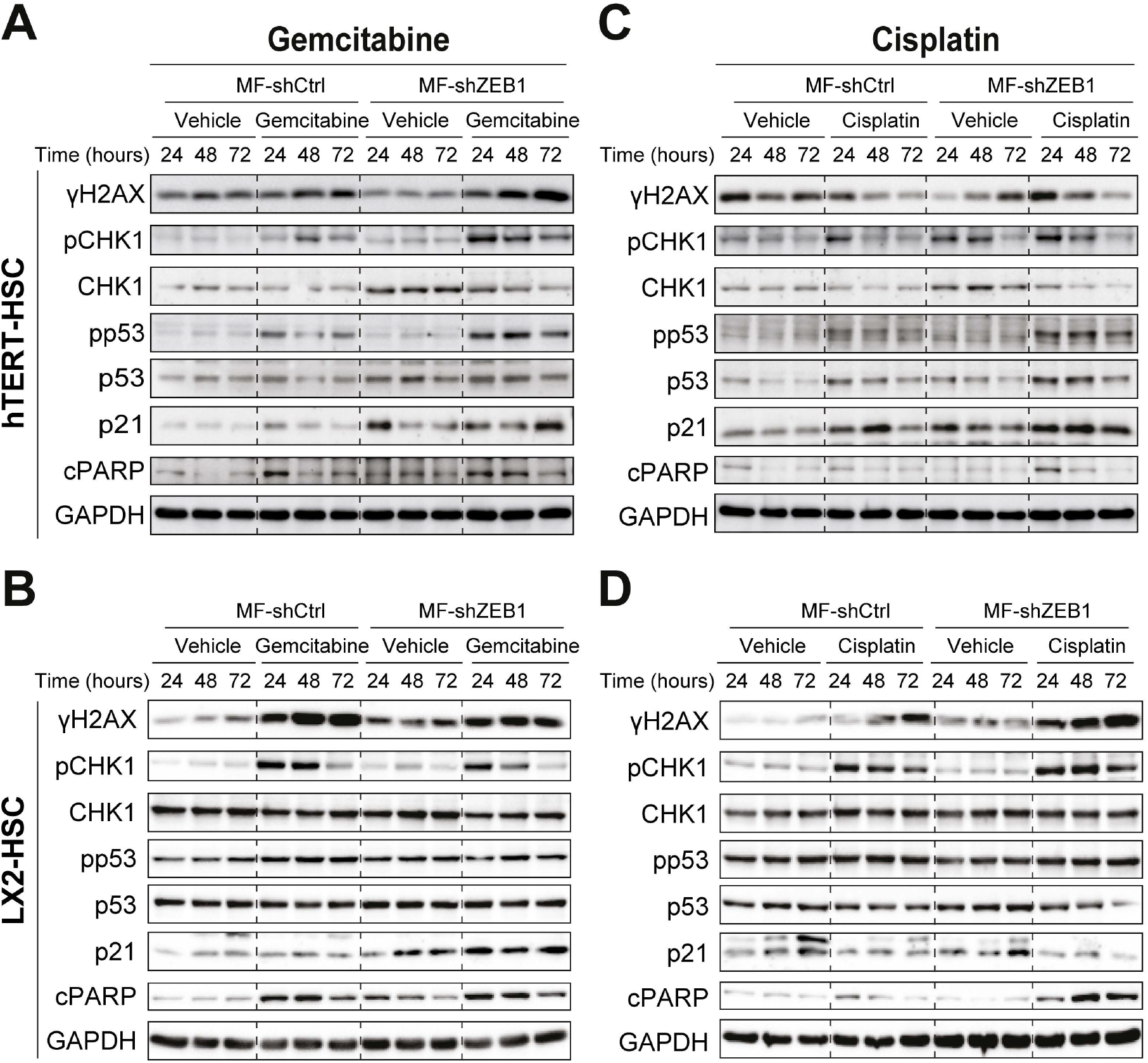
ZEB1 reduces apoptosis induced by gemcitabine and cisplatin in liver myofibroblasts. Representative images of Western blot analysis of phosphorylated H2AX, phosphorylated and total CHK1, phosphorylated and total p53, p21 and cleaved PARP (cPARP) in shRNA-Control (shCtrl) and shRNA-ZEB1 hTERT-HSC and LX2-HSC cells treated with gemcitabine (**A-B**) or cisplatin (**C-D**).

### 3.3. ZEB1 down-regulation sensitizes CAF to gemcitabine- and cisplatin-induced apoptosis

Gemcitabine and cisplatin are genotoxic drugs known to induce DNA damage, triggering intracellular signalling pathways that lead to cell cycle arrest and ultimately death by apoptosis.[19] Immunoblot analysis revealed that ZEB1-downregulated CAF exposed to gemcitabine and cisplatin showed indeed a stronger signal for γH2AX and phosphorylation of CHK1 (except for LX2-HSC cells treated with gemcitabine), both critical regulators of the DNA damage response (Figure 3A-D). Similarly, shZEB1 cells exposed to chemotherapeutic drugs showed more p53 phosphorylation, p21 expression and cleaved of PARP, all indicative of cell cycle arrest and death by apoptosis (Figure 3A-D). Interestingly, these changes were less intense in LX2-HSC shZEB1 cells exposed to gemcitabine, consistent with the viability results shown in Figure 2B-C. This data indicates that ZEB1 downregulation enhances CAF sensitivity to apoptosis induced by genotoxic chemotherapeutic drugs.

### 3.4. ZEB1 down-regulation alters the anti/proapoptotic balance in liver myofibroblasts

The next step in our study was to decipher the mechanisms behind the chemoresistance (MOC) regulated by ZEB1 that renders CAF more resistant to chemotherapeutic drugs. One of the most common MOC is the alteration of transporters of these drugs.[20] The expression analysis of the main uptake transporters and export pumps of gemcitabine and cisplatin did not show any significant changes that could explain our previous results (Supplementary Figure 4A-B). Thus, given the sensitization to apoptotic death observed in ZEB1-downregulated cells, we decided to explore if the expression of genes involved in the regulation of apoptosis was modified in shZEB1 cells. Indeed, analysis of expression of the BCL2 family members showed several changes that could account for the increased sensitivity of our cells (Figure 4A-B). Among those changes, two genes stand out among all. We observed a statistically significant downregulation of the antiapoptotic gene *BCL2*, along with an upregulation of the proapoptotic gene *BMF* in both shZEB1 cell lines (Figure 4A-B), indicating that reduced ZEB1 expression altered the anti/proapoptotic balance of the CAF.

**Figure 4.**
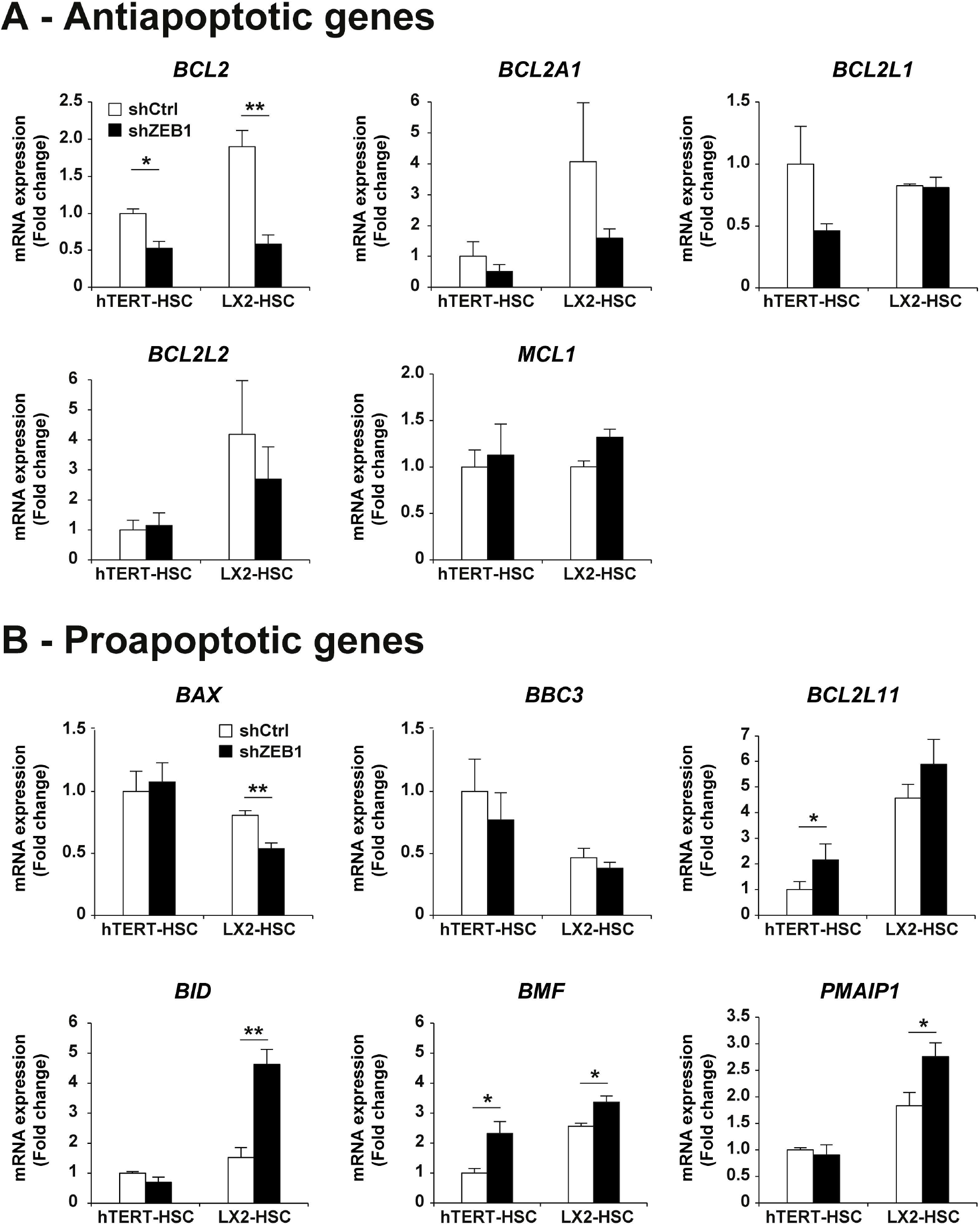
ZEB1 disrupts the pro-/anti-apoptotic balance in liver myofibroblasts. Changes in the mRNA expression of (A) antiapoptotic and (B) proapoptotic genes from the BCL2 superfamily in shRNA-Control (shCtrl) and shRNA-ZEB1 hTERT-HSC and LX2-HSC cells. Results are expressed as means±SEM from 5 independent cultures. *, p < 0.05; **, p < 0.01; as compared with shCtrl cells.

To understand the importance of this relationship of *ZEB1* with *BCL2* and *BMF* we analysed their expression and correlation in human iCCA samples. Interestingly, *ZEB1* expression correlated with *BCL2* expression in samples obtained by laser microdissection of iCCA stroma, but not in samples from the whole tumour from two independent cohorts (Figure 5A), indicating that, as our previous results suggested, ZEB1 role regulating the anti/proapoptotic balance and the sensitivity to chemotherapeutic drugs is restricted to the TME, but not to tumour cells. On the contrary, *BMF* did not significantly correlate with *ZEB1* in any of the cohorts analysed (Supplementary Figure 5). Thus, we focused the further investigation in the relationship between *ZEB1* and *BCL2*. Interestingly, a recent study determined by chip-seq ZEB1 binding sites in melanoma cells, where ZEB1 plays a key role as a major transcriptional regulator of cell state transitions.[15] We further analysed the data from this study to show that ZEB1-expressing cells bind to the promoter of BCL2 in a region located between 63317981-63318498 from chromosome 18 (Figure 5C). Interestingly, further interrogation of this region showed 5 E-BOX binding motifs with the consensus sequence for ZEB1 binding (CACCTG) (Figure 5C). Altogether, these results suggest that ZEB1 directly regulates *BCL2* expression in liver myofibroblasts.

**Figure 5.**
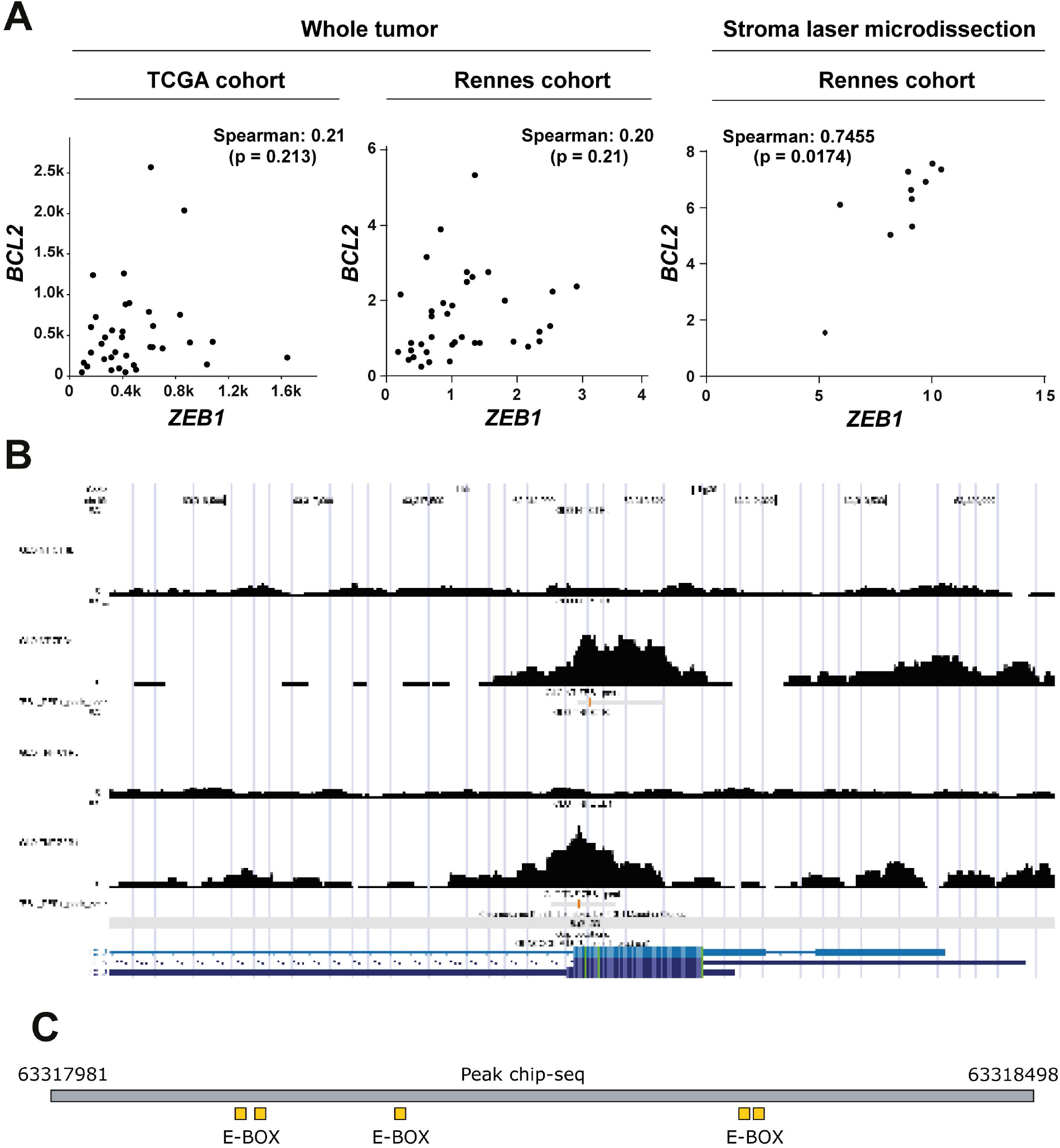
ZEB1 regulates BCL2 expression. **A**. Correlation between *ZEB1* and *BCL2* expression in two independent cohorts of human cholangiocarcinoma from whole tissue (TCGA and Rennes cohorts) and in 10 tumor stromal samples from the Rennes Cohort isolated by laser microdissection. **B**. ChIP-PCR analysis from GSE246672 indicating ZEB1 binding peaks on *BCL2* gene in ZEB1 expressing cells (GLO non-treated (NT) and TNF-α treated cells). **C**. Scheme showing E-BOX elements with ZEB1 consensus binding sequence in the genomic region of *ZEB1* peak from B (63317981-63318498).

### 3.5. BCL2 regulates the sensitivity to gemcitabine and cisplatin in CAF

To further understand the functional relationship between ZEB1 and BCL2 in chemoresistance and its potential therapeutic use, we first employed a specific inhibitor of BCL2, called venetoclax, on hTERT-HSC shControl cells. As shown on Figure 6A, venetoclax sensitized shControl cells to both gemcitabine and cisplatin, with a stronger effect observed for cisplatin. Venetoclax reduced cell viability (Figure 6A) and increased the cleavage of PARP (Figure 6B) leading to cell death, easily visualized by transmitted light microscopy (Figure 6C). Then, in order to perform a mirror experiment, we generated a cell line derived from hTERT-HSC shZEB1 cells that overexpress BCL2 (BCL2ov) (Supplementary Figure 6A-C). As expected, BCL2 overexpression partially protected hTERT-shZEB1 cells to both gemcitabine and cisplatin (Figure 6D). Accordingly, we observed reduced cleavage of PARP after exposure to both drugs in BCL2 overexpressing cells (Figure 6E), as well as more viable cells at the microscope (Figure 6F). Therefore, BCL2 regulates chemoresistance of CAF to gemcitabine and cisplatin.

**Figure 6.**
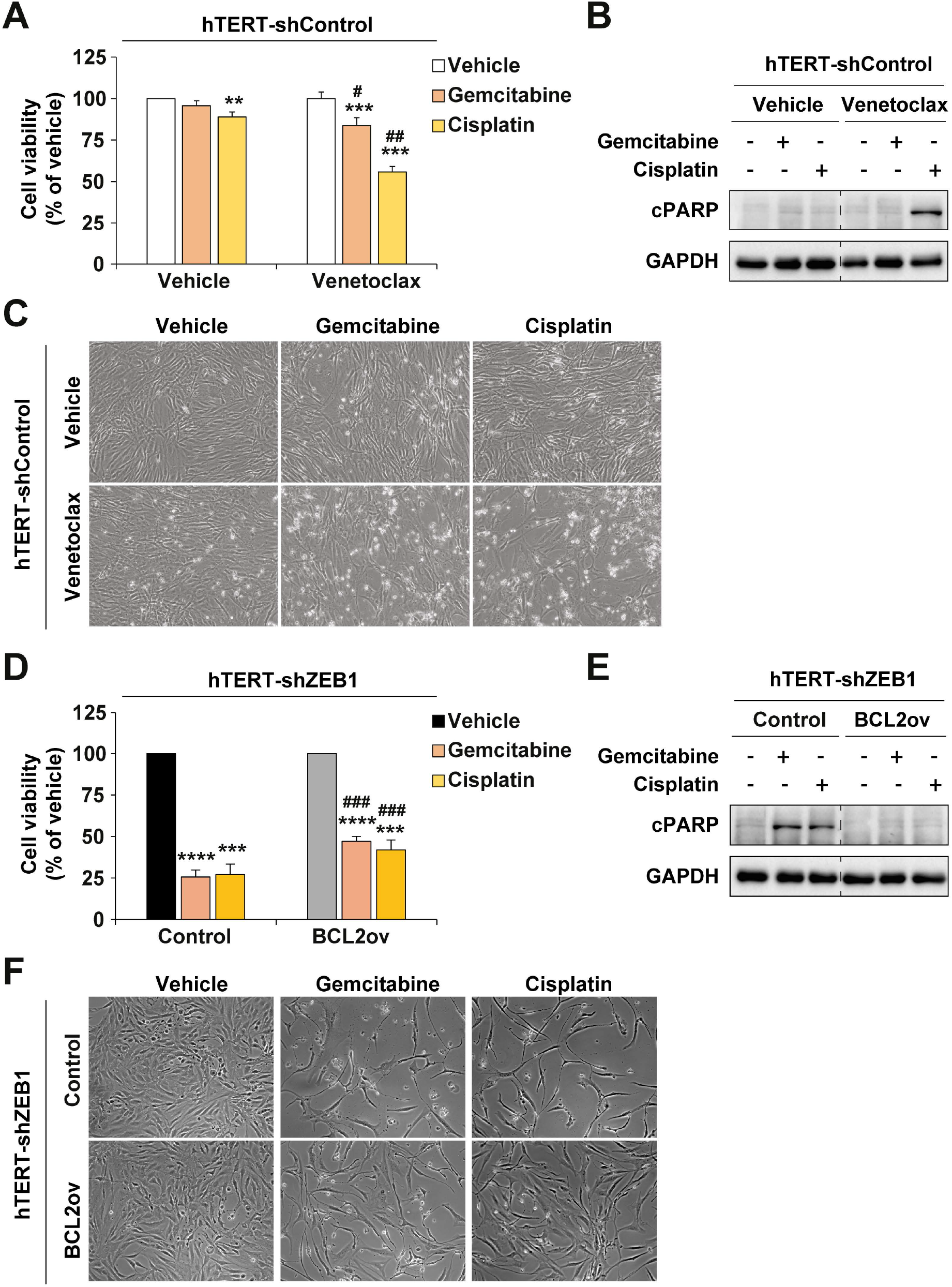
BCL2 induces chemoresistance to gemcitabine and cisplatin in liver myofibroblasts. **A**. Effect of venetoclax (5 µM) on the response to gemcitabine (0.1 µM) and cisplatin (25 µM) of hTERT shRNA-Control cells. Cell viability was measured by the crystal violet test after incubation with the indicated drugs for 72h. **B**. Representative images of Western blot analysis of cleaved PARP in cells from A after 24 h of incubation. **C**. Representative pictures of cells from A at term. **D**. Effect of BCL2 overexpression (BCL2ov) on the response to gemcitabine (0.1 µM) and cisplatin (25 µM) of hTERT shRNA-ZEB1 cells. **E**. Representative images of Western blot analysis of cleaved PARP in cells from D after 24 h of incubation. **F**. Representative pictures of cells from D at term. Results are expressed as means±SEM from 5 independent cultures. *, p < 0.05; **, p < 0.01; ***, p < 0.001; ****, p < 0.0001 as compared with vehicle. #, p < 0.05; ##, p < 0.01; ###, p < 0.001; as compared with hTERT-shRNA-Control-Vehicle or hTERT-shRNA-ZEB1-Control.

### 3.6. CAF with high ZEB1-BCL2 expression protect iCCA tumour cells from gemcitabine and cisplatin

To assess the potential contribution of ZEB1 expression in CAF to iCCA resistance to gemcitabine and cisplatin, we conducted spheroid studies. First, we determined the sensitivity of tumour spheroids formed solely by HuCCT1 tumour cells originating from iCCA by exposing them to the chemotherapeutic drugs and determining the spheroid size (Supplementary Figure 7A-C). Then, we developed mixed spheroids formed by HuCCT1 tumour cells in combination with either hTERT-HSC shControl or shZEB1 (Figure 7A) and exposed them to gemcitabine or cisplatin in absence or in presence of the BCL2 inhibitor venetoclax. First, we observed that the spheres formed by HuCCT1 cells + hTERT-HSC shZEB1 cells were smaller than those formed by HuCCT1 + hTERT-HSC shControl cells (Figure 7B and D). These data corroborate our previous studies performed with conditioned medium regarding the potential of hTERT-HSC to promote iCCA cells growth.[6] Second, the addition of venetoclax alone did not have an effect on the size of spheroids formed by HuCCT1 and hTERT-HSC shControl (Figure 7B and D). However, when the spheroids were exposed to gemcitabine or cisplatin, we could observe that 1/addition of shControl CAF to the spheres increases the chemoresistance of the spheroids compared to tumour cells alone (Supplementary Figure 7B-C and Figure 7C-D), 2/ that spheres formed by HuCCT1 cells + hTERT-HSC shZEB1 were more sensitive than those that contained shControl cells, and 3/the exposure to venetoclax increased the sensitivity of spheroids formed by HuCCT1 cells + hTERT-HSC shControl cells to levels similar to those that contain shZEB1 (Figure 7C-D). Following the previous experimental plan, we decided to use our hTERT-HSC BCL2ov to perform a mirrored experiment. When we exposed mixed spheroids formed by HuCCT1 cells + hTERT-HSC shZEB1-Control or BCL2ov to gemcitabine and cisplatin, we could observe that BCL2 overexpression in hTERT-HSC cells does not impact the growth of the spheroids in absence of drugs (Figure 7E and 7G), but it protected them from the toxicity of gemcitabine and cisplatin (Figure 7F-G). These results indicate that BCL2 inhibition mimics the effect of ZEB1 downregulation in CAF model, sensitizing the tumour cells to chemotherapeutic drugs and confirming the link between ZEB1 and BCL2 in this process.

**Figure 7.**
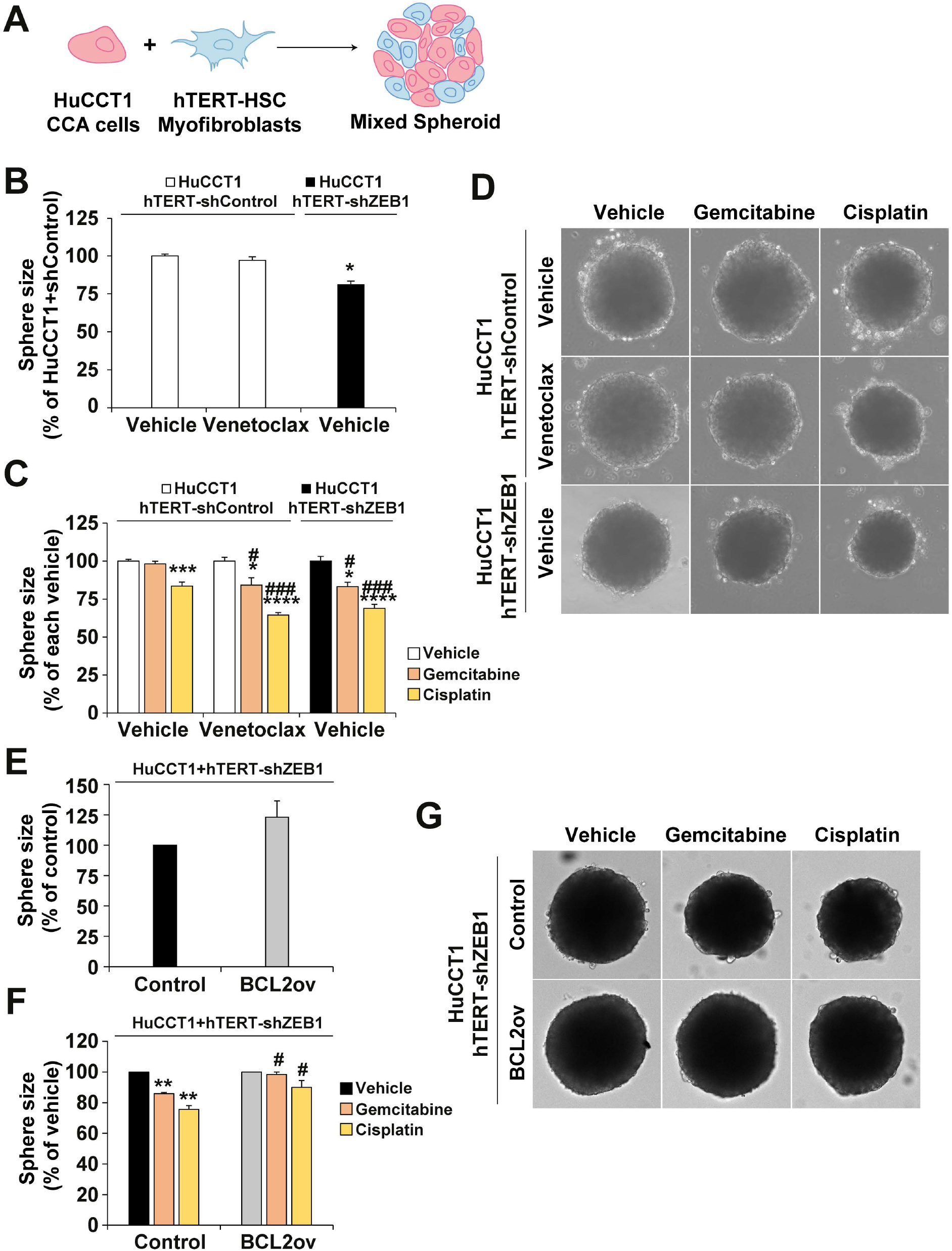
BCL2 from liver myofibroblasts induce chemoresistance to gemcitabine and cisplatin in 3D models of intrahepatic cholangiocarcinoma (iCCA). **A**. Scheme depicting the mixed spheroids used in the experiments. **B**. Size of spheroids containing HuCCT1 cells plus hTERT-shRNA-Ctrl or hTERT-shRNA-ZEB1 in presence of the vehicle or venetoclax. **C**. Effect of venetoclax on the sensibility of mixed spheroids to gemcitabine and cisplatin. **D**. Representative images of the spheres quantified in B and C. **E**. Size of spheroids containing HuCCT1 cells plus hTERT-shZEB1-Control o hTERT-shZEB1-BCL2ov. **F**. Effect of BCL2 overexpression (BCL2ov) in hTERT-shZEB1 cells on the sensibility of mixed spheroids to gemcitabine and cisplatin. **G**. Representative images of the spheres quantified in F and G. Results are expressed as means±SEM from 5 independent cultures. *, p < 0.05; **, p < 0.01; ***, p < 0.001; ****, p < 0.0001 as compared with vehicle. #, p < 0.05; ##, p < 0.01; ###, p < 0.001; as compared with HuCCT1+hTERT-shCtrl-Vehicle or HuCCT1+hTERT-shZEB1-Control.

## 4. Discusion

ZEB1 was originally identified as an EMT-TF involved in embryonic development.[21] Since then it has been involved in different processes related to carcinogenesis, metastasis and chemoresistance to several drugs, including chemotherapies and targeted therapies.[7-9, 22, 23]. Recently, we have described a role for ZEB1 in the acquisition of a more aggressive phenotype of iCCA tumour cells.[6] Furthermore, we showed that ZEB1 is involved in different aspects of iCCA progression through its expression in the tumour microenvironment, more specifically in CAF where ZEB1 regulates the activation status of the cells and controls their crosstalk through soluble factors with iCCA tumour cells.[6] However, the information about the role of ZEB1 in CAF is still scarce and, thus, we followed up our previous work in the present study. Here we have demonstrated that ZEB1 is expressed in CAF from iCCA in virtually all patients independently of the tumour grade. Further sc-RNAseq analysis showed that CAF is the population with the highest ZEB1 expression in the TME of iCCA, indicating the importance of this EMT-TF in CAF biology and iCCA pathology.

The recent approval of durvalumab and pembrolizumab for the treatment of iCCA patients represents the first change in the standard of care for the majority of iCCA patients in the last decade.[3, 4] Nevertheless, suboptimal responses in an important percentage of patients underlies the necessity of further investigation to improve current therapies for iCCA patients. Furthermore, these immunotherapeutic drugs are still given in combination with the traditional chemotherapeutic combination of gemcitabine and cisplatin that iCCA patients receive. Thus, research focused on understanding the MOC to gemcitabine and cisplatin are still of great interest.[20] In this sense, CAF have been related to chemoresistance to gemcitabine and cisplatin in other cancers,[24-26] which led us to test this possibility in iCCA. Indeed, our experiments on mixed spheroids formed by iCCA cells and liver myofibroblasts show that the presence of the later in the spheroids protects tumour cells against the toxicity of chemotherapeutic drugs.

Upon the above information we wanted to evaluate the possibility of ZEB1 expression in CAF playing a role in iCCA chemoresistance to present therapies. Interestingly, we found that fibroblasts depleted for ZEB1 (shRNA-ZEB1) showed higher sensitivity to gemcitabine and cisplatin, but not to sorafenib, indicating ZEB1 protects these cells specifically against iCCA therapeutic regimens, but not HCC targeted therapies. While several reports show that ZEB1 protects tumour cells to different drugs,[9, 27] to our knowledge, this is the first report showing that ZEB1 is able to induce chemoresistance in fibroblasts. Furthermore, when we interrogated the mechanisms responsible for this chemoresistance we observed surprisingly that one of the main MOC involved in the resistance to these drugs, the modification of their transport across the membrane by different uptake and export pumps,[20] was unchanged. However, we observed that ZEB1 protected liver myofibroblasts from gemcitabine- and cisplatin-induced apoptosis via regulation of BCL2, which we observed both in cell lines and in human iCCA in sample from microdissected stroma. The relationship between ZEB1 and BCL2 expression has been previously stressed in mantle cell lymphoma cells. In these cells downregulation of ZEB1 also led to downregulation of BCL2, as in our fibroblasts, and also led to increased sensitivity to different chemotherapeutics including gemcitabine.[28] Here we show the specific region where ZEB1 binds *BCL2* regulatory regions to regulate the expression of BCL2 thanks to analysis of public data from a recent study.[15]

Our data open the way for the potential therapeutic usefulness of targeting ZEB1 in late stages of tumour development to sensitize iCCA to current therapeutic regimens. Indeed, our discovery of BCL2 regulation by ZEB1 opened the door to target ZEB1 functions through the use of BCL2 inhibitors. Indeed, venetoclax, a specific BCL2 inhibitor currently used in clinical trials, was able to sensitize both liver myofibroblasts and mixed spheroids to gemcitabine and cisplatin to the levels of cells depleted from ZEB1. Mirrored experiments with shRNA-ZEB1 overexpressing BCL2 confirmed the prominent role of ZEB1-BCL2 axis in the chemoresistance of these cells to the chemotherapeutic drugs. Previous studies show that depletion of CAF with similar molecules, such as navitoclax, reduced iCCA progression.[29]

A question that remains unanswered is how the resistant fibroblasts that express ZEB1 are also able to provide resistance to iCCA cells against gemcitabine and cisplatin. We have previously shown evidence that ZEB1 is at the interface of tumour–stroma cross-communication by regulating the expression of growth factors and proinflammatory cytokines that impact iCCA progression. Thus, perhaps the greater presence of fibroblasts producing growth factors provides iCCA cells with enough support to continue proliferating and growing. Fuerthermore, a recent report indicates that IL8 derived from CAF induces cisplatin resistance in lung cancer,[25] and we showed in our previous study that IL8 may be regulated by ZEB1 also in liver myofibroblasts. This suggest that there may exist additional mechanisms regulated by ZEB1 involved in iCCA chemoresistance. Further trnacriptomic and proteomic analyses will surely open new avenues for understanding ZEB1 roles in iCCA TME and ensure further research on this topic in the coming years.

## 5- Conclusion

This study underlines once again the importance of the roles played by ZEB1 in CAF from iCCA. We demonstrate that ZEB1 disrupts the pro/anti-apoptotic balance through the regulation of BCL2 in these cells and induces chemoresistance to gemcitabine and cisplatin. Furthermore, use of BCL2 inhibitors could represent an important tool to sensitize iCCA to the current chemotherapeutic drugs used in the vast majority of the patients.

## Abbreviations

(α-SMA): Alpha-smooth muscle actin
(CAF): cancer associated fibroblasts
(ChIP-PCR): Chromatin immunoprecipitation PCR
(EMT-TF): EMT-inducing transcription factor
(ECM): extracellular matrix
(HSC): hepatic stellate cell
(HCC): hepatocellular carcinoma
(iCCA): intrahepatic cholangiocarcinoma
(MOC): mechanism of chemoresistance
(shRNAs): small hairpin RNAs
(TMA): Tissue MicroArray
(TME): tumour microenvironment.

## CRediT authorship contribution statement

**Ester Gonzalez-Sanchez**: Data curation, Review & editing. **Marie Vallette**: Data curation, Review & editing. **Allan Pavy**: Data curation, Review & editing. **Aashreya Ravichandra**: Data curation, Review & editing. **Mirko Minini**: Data curation, Review & editing. **Josep Amengual**: Data curation, Review & editing. **Itziar Rodríguez-Rivas**: Data curation, Review & editing. **Carlos Andres Roldán-Hernández**: Data curation, Review & editing. **Corentin Louis**: Data curation, Review & editing. **Isabel Fabregat**: Data curation, Review & editing. **Nathalie Guedj**: Data curation, Review & editing. **Valérie Paradis**: Data curation, Review & editing. **Cedric Coulouarn**: Data curation, Review & editing. **Lynda Aoudjehane**: Data curation, Review & editing. **Laura Fouassier**: Data curation, Supervision, Resources, Project administration, Funding acquisition, Review & editing. **Javier Vaquero**: Data curation, Supervision, Resources, Project administration, Funding acquisition Conceptualization, Writing – original draft – review & editing.

## Data availability

All data are available in the manuscript.

## Acknowledgments

The authors acknowledge the common services from CIC-IBMCC in particular the Central Store Room coordinated by Sonia Perez Diaz and Maria Eugenia Fernandez de la Torre, and the Microscopy Unit coordinated by Ana Isabel Garcia. We thank Profs Dominique Wendum and Magali Svrcek from the Pathology Department, Prof François Paye from Digestive Surgical Surgery Departments, and Dr. Marie Lequoy from the Hepatology Department of Saint-Antoine Hospital, Assistance Publique – Hôpitaux de Paris (Direction de la Recherche Clinique et de l’Innovation), France, France. This article is based upon work from the COST Action CA22125 Precision medicine in biliary tract cancer (Precision-BTC-Network) supported by COST (European Cooperation in Science and Technology: www.cost.eu), in collaboration with the European Network for the Study of Cholangiocarcinoma (ENS-CCA: http://www.enscca.org/).

## Appendix A. Supplementary material

Supplementary data associated with this article can be found in the online version.

## References

1. Banales, J. M., Cardinale, V., Carpino, G., Marzioni, M., Andersen, J. B., Invernizzi, P., Lind, G. E., Folseraas, T., Forbes, S. J., Fouassier, L., Geier, A., Calvisi, D. F., Mertens, J. C., Trauner, M., Benedetti, A., Maroni, L., Vaquero, J., Macias, R. I., Raggi, C., Perugorria, M. J., Gaudio, E., Boberg, K. M., Marin, J. J. & Alvaro, D. (2016) Expert consensus document: Cholangiocarcinoma: current knowledge and future perspectives consensus statement from the European Network for the Study of Cholangiocarcinoma (ENS-CCA), Nat Rev Gastroenterol Hepatol. 13, 261–80.

2. Banales, J. M., Marin, J. J. G., Lamarca, A., Rodrigues, P. M., Khan, S. A., Roberts, L. R., Cardinale, V., Carpino, G., Andersen, J. B., Braconi, C., Calvisi, D. F., Perugorria, M. J., Fabris, L., Boulter, L., Macias, R. I. R., Gaudio, E., Alvaro, D., Gradilone, S. A., Strazzabosco, M., Marzioni, M., Coulouarn, C., Fouassier, L., Raggi, C., Invernizzi, P., Mertens, J. C., Moncsek, A., Ilyas, S. I., Heimbach, J., Koerkamp, B. G., Bruix, J., Forner, A., Bridgewater, J., Valle, J. W. & Gores, G. J. (2020) Cholangiocarcinoma 2020: the next horizon in mechanisms and management, Nat Rev Gastroenterol Hepatol. 17, 557–588.

3. Burris, H. A., 3rd, Okusaka, T., Vogel, A., Lee, M. A., Takahashi, H., Breder, V., Blanc, J. F., Li, J., Bachini, M., Zotkiewicz, M., Abraham, J., Patel, N., Wang, J., Ali, M., Rokutanda, N., Cohen, G. & Oh, D. Y. (2024) Durvalumab plus gemcitabine and cisplatin in advanced biliary tract cancer (TOPAZ-1): patient-reported outcomes from a randomised, double-blind, placebo-controlled, phase 3 trial, Lancet Oncol. 25, 626–635.

4. Kelley, R. K., Ueno, M., Yoo, C., Finn, R. S., Furuse, J., Ren, Z., Yau, T., Klumpen, H. J., Chan, S. L., Ozaka, M., Verslype, C., Bouattour, M., Park, J. O., Barajas, O., Pelzer, U., Valle, J. W., Yu, L., Malhotra, U., Siegel, A. B., Edeline, J., Vogel, A. & Investigators, K.-. (2023) Pembrolizumab in combination with gemcitabine and cisplatin compared with gemcitabine and cisplatin alone for patients with advanced biliary tract cancer (KEYNOTE-966): a randomised, double-blind, placebo-controlled, phase 3 trial, Lancet. 401, 1853–1865.

5. Cantallops Vila, P., Ravichandra, A., Agirre Lizaso, A., Perugorria, M. J. & Affo, S. (2024) Heterogeneity, crosstalk, and targeting of cancer-associated fibroblasts in cholangiocarcinoma, Hepatology. 79, 941–958.

6. Lobe, C., Vallette, M., Arbelaiz, A., Gonzalez-Sanchez, E., Izquierdo, L., Pellat, A., Guedj, N., Louis, C., Paradis, V., Banales, J. M., Coulouarn, C., Housset, C., Vaquero, J. & Fouassier, L. (2021) Zinc Finger E-Box Binding Homeobox 1 Promotes Cholangiocarcinoma Progression Through Tumor Dedifferentiation and Tumor-Stroma Paracrine Signaling, Hepatology. 74, 3194–3212.

7. Li, J., Kou, Y., Zhang, X., Xiao, X., Ou, Y., Cao, L., Guo, M., Qi, C., Wang, Z., Liu, Y., Shuai, Q., Wang, H. & Yang, S. (2022) Biochanin A inhibits lung adenocarcinoma progression by targeting ZEB1, Discov Oncol. 13, 138.

8. Lima de Oliveira, J., More Milan, T., Longo Bighetti-Trevisan, R., Fernandes, R. R., Machado Leopoldino, A. & Oliveira de Almeida, L. (2023) Epithelial-mesenchymal transition and cancer stem cells: A route to acquired cisplatin resistance through epigenetics in HNSCC, Oral Dis. 29, 1991–2005.

9. Zhou, Y., Zhou, Y., Wang, K., Li, T., Zhang, M., Yang, Y., Wang, R. & Hu, R. (2019) ROCK2 Confers Acquired Gemcitabine Resistance in Pancreatic Cancer Cells by Upregulating Transcription Factor ZEB1, Cancers (Basel). 11.

10. Cerami, E., Gao, J., Dogrusoz, U., Gross, B. E., Sumer, S. O., Aksoy, B. A., Jacobsen, A., Byrne, C. J., Heuer, M. L., Larsson, E., Antipin, Y., Reva, B., Goldberg, A. P., Sander, C. & Schultz, N. (2012) The cBio cancer genomics portal: an open platform for exploring multidimensional cancer genomics data, Cancer Discov. 2, 401–4.

11. Gao, J., Aksoy, B. A., Dogrusoz, U., Dresdner, G., Gross, B., Sumer, S. O., Sun, Y., Jacobsen, A., Sinha, R., Larsson, E., Cerami, E., Sander, C. & Schultz, N. (2013) Integrative analysis of complex cancer genomics and clinical profiles using the cBioPortal, Sci Signal. 6, pl1.

12. Angenard, G., Merdrignac, A., Louis, C., Edeline, J. & Coulouarn, C. (2019) Expression of long non-coding RNA ANRIL predicts a poor prognosis in intrahepatic cholangiocarcinoma, Dig Liver Dis. 51, 1337–1343.

13. Sulpice, L., Rayar, M., Desille, M., Turlin, B., Fautrel, A., Boucher, E., Llamas-Gutierrez, F., Meunier, B., Boudjema, K., Clement, B. & Coulouarn, C. (2013) Molecular profiling of stroma identifies osteopontin as an independent predictor of poor prognosis in intrahepatic cholangiocarcinoma, Hepatology. 58, 1992–2000.

14. Vaquero, J., Lobe, C., Tahraoui, S., Claperon, A., Mergey, M., Merabtene, F., Wendum, D., Coulouarn, C., Housset, C., Desbois-Mouthon, C., Praz, F. & Fouassier, L. (2018) The IGF2/IR/IGF1R Pathway in Tumor Cells and Myofibroblasts Mediates Resistance to EGFR Inhibition in Cholangiocarcinoma, Clin Cancer Res. 24, 4282–4296.

15. Durand, S., Tang, Y., Pommier, R. M., Benboubker, V., Grimont, M., Boivin, F., Barbollat-Boutrand, L., Cumunel, E., Dupeuble, F., Eberhardt, A., Plaschka, M., Dalle, S. & Caramel, J. (2024) ZEB1 controls a lineage-specific transcriptional program essential for melanoma cell state transitions, Oncogene. 43, 1489–1505.

16. Barrett, T., Wilhite, S. E., Ledoux, P., Evangelista, C., Kim, I. F., Tomashevsky, M., Marshall, K. A., Phillippy, K. H., Sherman, P. M., Holko, M., Yefanov, A., Lee, H., Zhang, N., Robertson, C. L., Serova, N., Davis, S. & Soboleva, A. (2013) NCBI GEO: archive for functional genomics data sets--update, Nucleic Acids Res. 41, D991–5.

17. Perez, G., Barber, G. P., Benet-Pages, A., Casper, J., Clawson, H., Diekhans, M., Fischer, C., Gonzalez, J. N., Hinrichs, A. S., Lee, C. M., Nassar, L. R., Raney, B. J., Speir, M. L., van Baren, M. J., Vaske, C. J., Haussler, D., Kent, W. J. & Haeussler, M. (2025) The UCSC Genome Browser database: 2025 update, Nucleic Acids Res. 53, D1243–D1249.

18. Vogel, A., Bridgewater, J., Edeline, J., Kelley, R. K., Klumpen, H. J., Malka, D., Primrose, J. N., Rimassa, L., Stenzinger, A., Valle, J. W., Ducreux, M. &, E. G. C. E. a. (2023) Biliary tract cancer: ESMO Clinical Practice Guideline for diagnosis, treatment and follow-up, Ann Oncol. 34, 127–140.

19. Gielecinska, A., Kciuk, M., Yahya, E. B., Ainane, T., Mujwar, S. & Kontek, R. (2023) Apoptosis, necroptosis, and pyroptosis as alternative cell death pathways induced by chemotherapeutic agents?, Biochim Biophys Acta Rev Cancer. 1878, 189024.

20. Marin, J. J. G., Lozano, E., Briz, O., Al-Abdulla, R., Serrano, M. A. & Macias, R. I. R. (2017) Molecular Bases of Chemoresistance in Cholangiocarcinoma, Curr Drug Targets. 18, 889–900.

21. Vandewalle, C., Van Roy, F. & Berx, G. (2009) The role of the ZEB family of transcription factors in development and disease, Cell Mol Life Sci. 66, 773–87.

22. Caramel, J., Ligier, M. & Puisieux, A. (2018) Pleiotropic Roles for ZEB1 in Cancer, Cancer Res. 78, 30–35.

23. Gohlke, L., Alahdab, A., Oberhofer, A., Worf, K., Holdenrieder, S., Michaelis, M., Cinatl, J., Jr. & Ritter, C. A. (2023) Loss of Key EMT-Regulating miRNAs Highlight the Role of ZEB1 in EGFR Tyrosine Kinase Inhibitor-Resistant NSCLC, Int J Mol Sci. 24.

24. Qi, R., Bai, Y., Li, K., Liu, N., Xu, Y., Dal, E., Wang, Y., Lin, R., Wang, H., Liu, Z., Li, X., Wang, X. & Shi, B. (2023) Cancer-associated fibroblasts suppress ferroptosis and induce gemcitabine resistance in pancreatic cancer cells by secreting exosome-derived ACSL4-targeting miRNAs, Drug Resist Updat. 68, 100960.

25. Wu, J., Zhang, Q., Wu, J., Yang, Z., Liu, X., Lou, C., Wang, X., Peng, J., Zhang, J., Shang, Z., Xiao, J., Wang, N., Zhang, R., Zhou, J., Wang, Y., Hu, Z., Zhang, R., Zhang, J. & Zeng, Z. (2024) IL-8 from CD248-expressing cancer-associated fibroblasts generates cisplatin resistance in non-small cell lung cancer, J Cell Mol Med. 28, e18185.

26. Che, Y., Wang, J., Li, Y., Lu, Z., Huang, J., Sun, S., Mao, S., Lei, Y., Zang, R., Sun, N. & He, J. (2018) Cisplatin-activated PAI-1 secretion in the cancer-associated fibroblasts with paracrine effects promoting esophageal squamous cell carcinoma progression and causing chemoresistance, Cell Death Dis. 9, 759.

27. Cui, Y., Qin, L., Tian, D., Wang, T., Fan, L., Zhang, P. & Wang, Z. (2018) ZEB1 Promotes Chemoresistance to Cisplatin in Ovarian Cancer Cells by Suppressing SLC3A2, Chemotherapy. 63, 262–271.

28. Sanchez-Tillo, E., Fanlo, L., Siles, L., Montes-Moreno, S., Moros, A., Chiva-Blanch, G., Estruch, R., Martinez, A., Colomer, D., Gyorffy, B., Roue, G. & Postigo, (2014) The EMT activator ZEB1 promotes tumor growth and determines differential response to chemotherapy in mantle cell lymphoma, Cell Death Differ. 21, 247–57.

29. Mertens, J. C., Fingas, C. D., Christensen, J. D., Smoot, R. L., Bronk, S. F., Werneburg, N. W., Gustafson, M. P., Dietz, A. B., Roberts, L. R., Sirica, A. E. & Gores, G. J. (2013) Therapeutic effects of deleting cancer-associated fibroblasts in cholangiocarcinoma, Cancer Res. 73, 897–907.

