## Supplementary Material for "ZEB1 regulates BCL2 in cancer-associated fibroblasts to promote cholangiocarcinoma chemoresistance to gemcitabine and cisplatin"

1-HepatoBiliary Tumours Lab, Centro de Investigación del Cáncer and Instituto de Biología Molecular y Celular del Cáncer, CSIC-Universidad de Salamanca, Salamanca 37007, Spain
2-Department of Physiology and Pharmacology, University of Salamanca, 37007, Salamanca, Spain
3- National Biomedical Research Institute on Liver and Gastrointestinal Diseases (CIBERehd), Instituto de Salud Carlos III, Madrid, Spain
4-Sorbonne Université, Inserm, Centre de Recherche Saint-Antoine, CRSA, F-75012 Paris, France.
5-TUM School of Medicine and Health, Department of Clinical Medicine – Clinical Department for Internal Medicine II, University Medical Center, Technical University of Munich, Germany.
6-Université Paris Cité, CNRS, Inserm, NABI, F-75006 Paris, France.
7-TGF-β and Cancer Group, Oncobell Program, Bellvitge Biomedical Research Institute (IDIBELL), Barcelona, Spain
8-Inserm, Univ Rennes, OSS (Oncogenesis, Stress, Signaling) UMR_S 1242, Centre de Lutte contre le Cancer Eugène Marquis, F-35042, Rennes, France
9-Bioinformatic Platform, CIBERehd, Barcelona, Spain.
10- Service d'Anatomie Pathologique Hôpital Beaujon, F-92110 Clichy, France; INSERM, UMR 1149, Centre de Recherche sur l'Inflammation, F-75018, Paris, France
11-ICAN BioCell-Human Liver Biology core, IHU - Foundation for innovation in Cardiometabolism and Nutrition (IHU-ICAN), F-75013 Paris, France.
12-Sorbonne Université, INSERM, UMRS 1166, F-75013, Paris, France.

* Share first authorship
### Share senior authorship

**Corresponding author:**

Laura Fouassier, NABI (Nanomédecine, Biologie Extracellulaire, Intégratome et Innovations en santé) CNRS UMR 8175, INSERM U1334, Université Paris Cité, 75006 Paris, France, CNRS UMR 8175, INSERM U1334. Phone: +33 (0)6 98 77 40 01,

Javier Vaquero, HepatoBiliary Tumours Lab, Centro de Investigación del Cáncer and Instituto de Biología Molecular y Celular del Cáncer, CSIC-Universidad de Salamanca, Salamanca 37007, Spain. Phone: +34 626569867,

**Supplementary material**

**Supplementary Table 1. Human primers used for quantitative real-time PCR.**

| Gene | Protein | Forward (5'→3') | Reverse (5'→3') |
|---|---|---|---|
| *ABCC1* | MRP1 | CCATGTGGGAAAACACATCTT | CTGTGCGTGACCAAGATCC |
| *ABCC2* | MRP2 | TGAAGAGGAAGCCACAGTCCATGA | TTCAGATGCCTGCCATTGGACCTA |
| *ABCC10* | ABCC10 | TTCATCACCTATGTCCTCATGG | AATGAGCATTCGCACCAGT |
| *ABCG2* | ABCG2 | CCCAGGCCTCTATAGCTCAGATCATT | CACGGCTGAAACACTGCTGAAACA |
| *ATP7A* | ATP7A | AAGAAATGATCAACCTTCATTCTTC | CCTCTGATGTTTTGCCCTGT |
| *ATP7B* | ATP7B | GATGCCTGAGCAGGAGAGAC | GGGTAGGCAAAGAAAGCTTAGA |
| *BAX* | BAX | CCCGAGAGGTCTTTTTCCGAG | CCAGCCCATGATGGTTCTGAT |
| *BBC3* | BBC3 | GACCTCAACGCACAGTACGA | GAGATTGTACAGGACCCTCCA |
| *BCL2* | BCL2 | GGTGGGGTCATGTGTGTGG | CGGTTCAGGTACTCAGTCATCC |
| *BCL2A1* | BCL2A1 | TTGCTCTCCACCAGGCAGAAGAT | GGACTGAGAACGCAACATTTTGTAGCAC |
| *BCL2L1* | BCL2L1 | CCTGCCTGCCTTTGCCTAA | CCCGGTTGCTCTGAGACATT |
| *BCL2L2* | BCL2L2 | TCACCCAGGTCTCCGATGAACT | GCTGTGGATCCAGTCAGCCAG |
| *BCL2L11* | BCL2L11 | TAAGTTCTGAGTGTGACCGAGA | GCTCTGTCTGTAGGGAGGTAGG |
| *BID* | BID | ATGGACCGTAGCATCCCTCC | GTAGGTGCGTAGGTTCTGGT |
| *BMF* | BMF | CAAATCTGAACAAGCCCAAGTCTTCCAG | CACACAGCTTAGTGAGCAGAACACAA |
| *GAPDH* | GAPDH | AGCCACATCGCTCAGACAC | GCCCAATACGACCAAATCC |
| *MCL1* | MCL1 | TGCTTCGGAAACTGGACATCA | TAGCCACAAAGGCACCAAAAG |
| *PMAIP1* | NOXA | GGAGATGCCTGGGAAGAAG | CCTGAGTTGAGTAGCACACTCG |
| *SLC29A1* | ENT1 | TGAGCGGAACTCTCTCAGTG | TGAGGTAGGTGAATAACAGCAGG |
| *SLC31A1* | CTR1 | TTCCTTCCCCATTCACATCT | ACAGAGTAAGGGGGCCAAAG |
| *ZEB1* | ZEB1 | ACAATCGTGGCCATTGCTGA | TGGTTCTTGGACTGCAGGGCT |

**Supplementary Table 2. Primary antibodies used for immunodetection.**

| Name | Species | Manufacturer | Reference | Dilution | Antigen unmasking |
|---|---|---|---|---|---|
| α-SMA | M | Dako | M851 | 1/500 (IHC)<br>1/100 (IF) | |
| BCL2 | M | Santa Cruz | Sc-509 | 1/1000 (WB) | |
| cPARP | R | CST | CST5625 | 1/1000 (WB) | |
| CHK1 | M | CST | CST2360 | 1/1000 (WB) | |
| pCHK1 | R | CST | CST2348 | 1/1000 (WB) | |
| GAPDH | M | Santa Cruz | sc-32233 | 1/5000 (WB) | |
| p21 | M | BD biosciences | 556431 | 1/500 (WB) | |
| p53 | M | Santa Cruz | sc-126 | 1/500 (WB) | |
| pp53 | R | CST | CST9284 | 1/1000 (WB) | |
| γH2A.X | R | CST | CST9718 | 1/1000 (WB) | |
| ZEB1 | R | Sigma | HPA027524 | 1/200 (IHC) | yes |
| ZEB1 | R | Santa Cruz | sc-25388 | 1/500 (WB)<br>1/50 (IF) | |

M, mouse; R, rabbit; WB, western blot; IF, immunofluorescence; IHC, immunohistochemistry.

**Supplementary Table 3. Clinical and pathological characteristics of patients with CCA (n=45)**

| | |
|---|---|
| Age (years)<br>Mean (± SD) | 62 +12 |
| Sex ratio (M/F) | 0.9 |
| Tumor size (mm)<br>Mean (± SD) | 82 (15-220) |
| Tumor grade | |
| Well differentiated | 17/45 (38%) |
| Moderately differentiated | 18/45 (40%) |
| Poorly differentiated | 10/45 (22%) |
| pTNM (7th edition) | |
| T1a | 13% |
| T1b | 9% |
| T2 | 78% |
| Vascular invasion | 71% |
| Perineural invasion | 31% |
| Lymph node metastasis | 37% |

**Supplementary Figures**

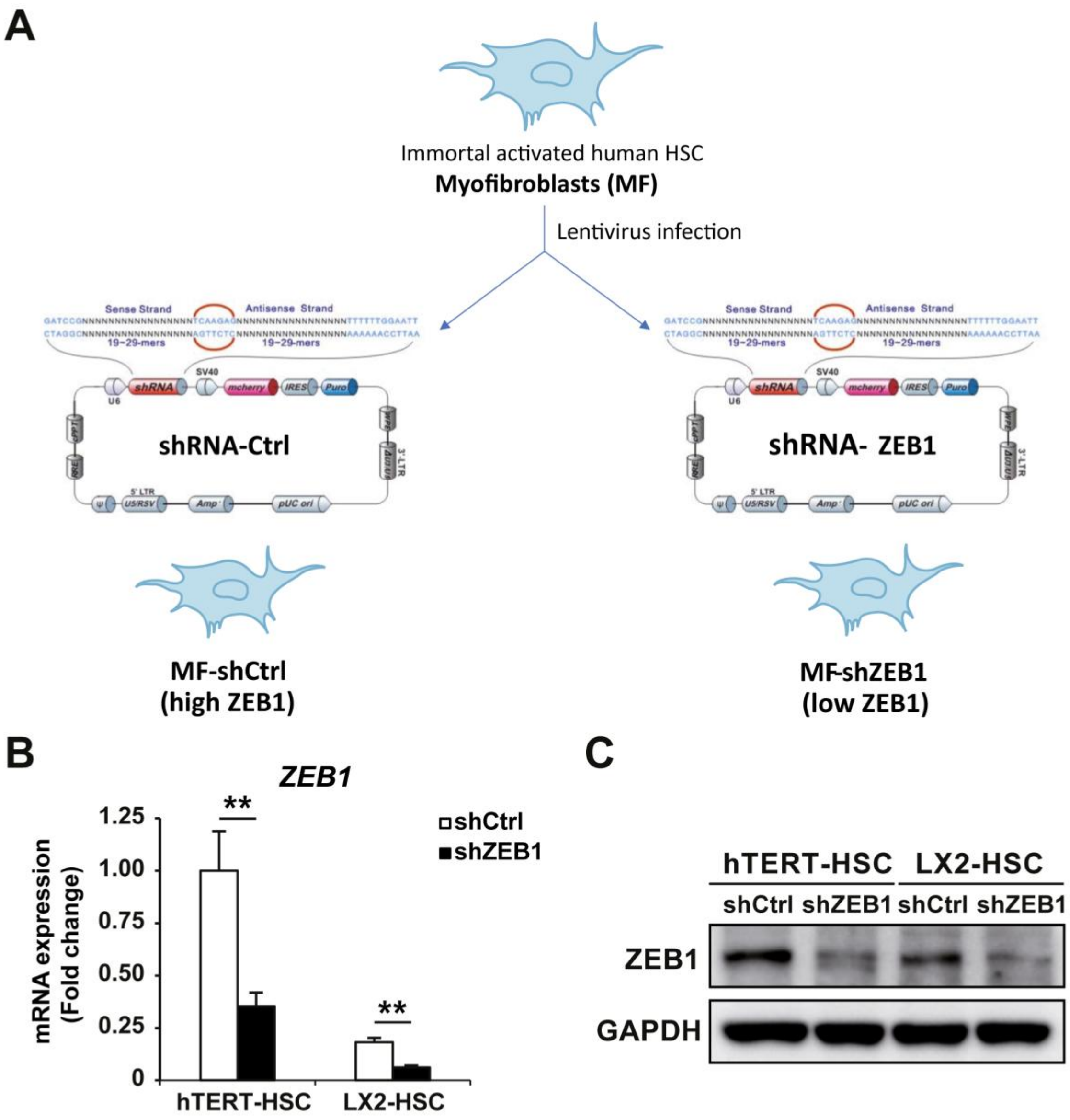


**Supplementary Figure 1. Generation of liver myofibroblasts with ZEB1 downregulation. A.** Scheme indicating the structure of the plasmids containing shRNA against ZEB1 used to produce the lentiviruses that were used to downregulate ZEB1. **B-C.** Changes in ZEB1 mRNA and protein expression in shRNA-Control (shCtrl) and shRNA-ZEB1 hTERT-HSC and LX2-HSC cells, determined by RT-QPCR and Western blot. Values are expressed as means ± SEM from 5 independent cultures. **, $p<0.01$

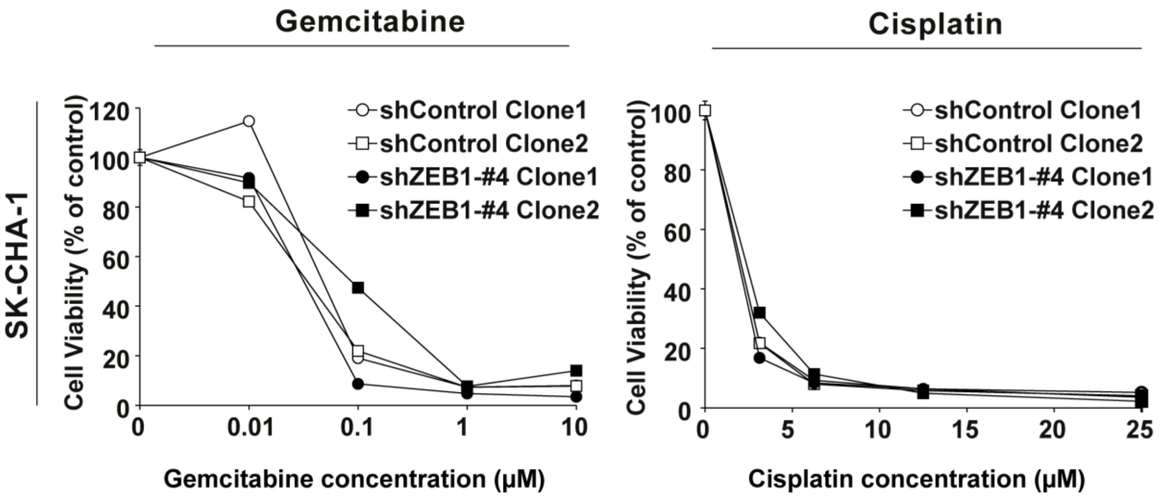


**Supplementary Figure 2. Downregulation of ZEB1 does not modify the sensitivity of cholangiocarcinoma (CCA) cells to gemcitabine or cisplatin.** Effect of gemcitabine and cisplatin on the viability of shRNA-Control (shControl) and shRNA-ZEB1 (shZEB1) SK-ChA-1 CCA cells. Cell viability was measured by the crystal violet test after incubation with the indicated concentrations of gemcitabine or cisplatin for 72h. Values are expressed as means ± SEM from at least 5 independent cultures.

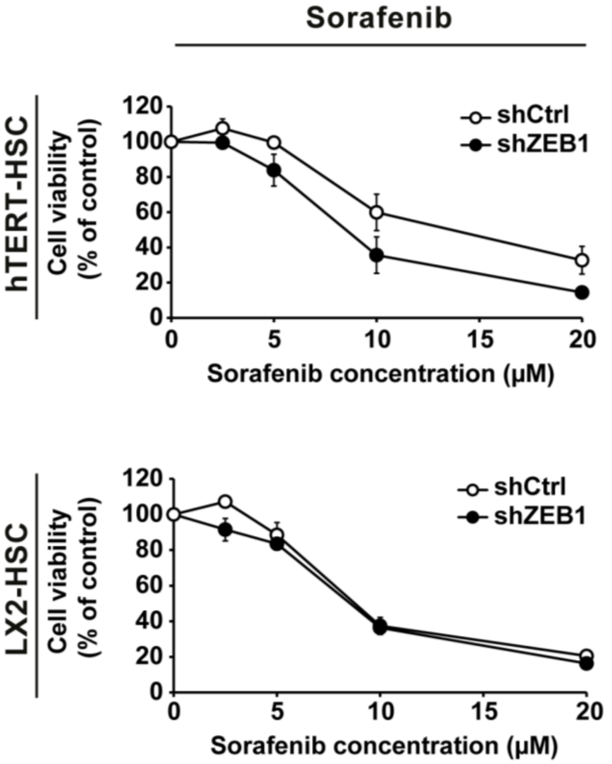


**Supplementary Figure 3. Downregulation of ZEB1 does not modify the sensitivity of liver myofibroblats to sorafenib.** Effect of sorafenib on the viability of shRNA-Control (shCtrl) and shRNA-ZEB1 hTERT-HSC and LX2-HSC cells. Cell viability was measured by the crystal violet test after incubation with the indicated concentrations of gemcitabine or cisplatin for 72h. Values are expressed as means ± SEM from at least 5 independent cultures.

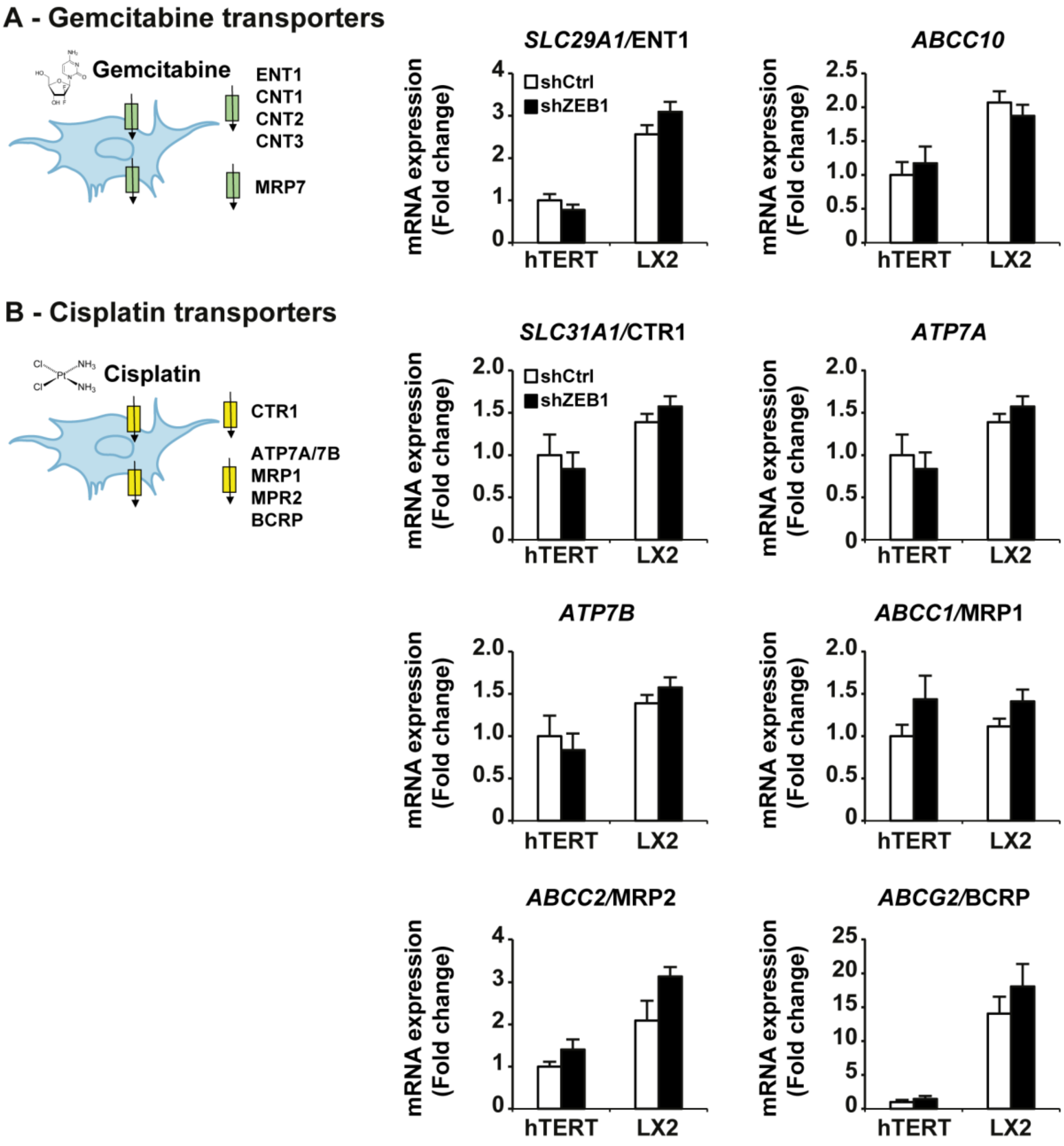


**Supplementary Figure 4. Downregulation of ZEB1 does not modify the expression of transporters from gemcitabine and cisplatin in liver myofibroblats.** Changes in the mRNA expression of uptake transporters and export pumps genes of gemcitabine (A) and cisplatin (B) in shRNA-Control (shCtrl) and shRNA-ZEB1 hTERT-HSC and LX2-HSC cells. Results are expressed as means ± SEM from 5 independent cultures.

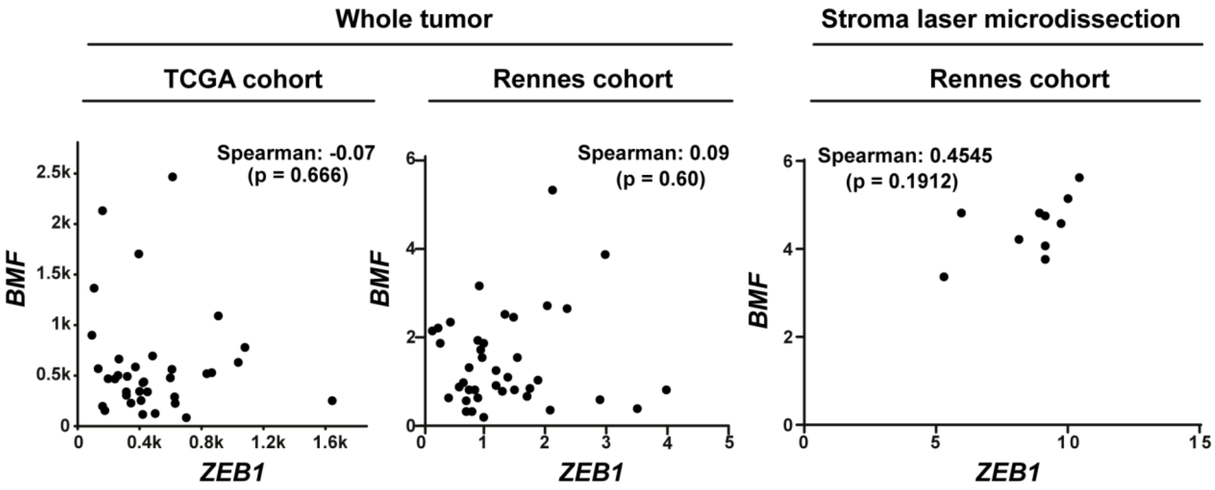


**Supplementary Figure 5. The expression of *ZEB1* does not correlate with that of *BMF* in human cholangiocarcinoma (CCA) samples. A.** Correlation between *ZEB1* and *BMF* expression in two independent cohorts of human CCA from whole tissue (TCGA and Rennes cohorts) or stroma extracted by laser microdissection (Rennes Cohort, n=10).

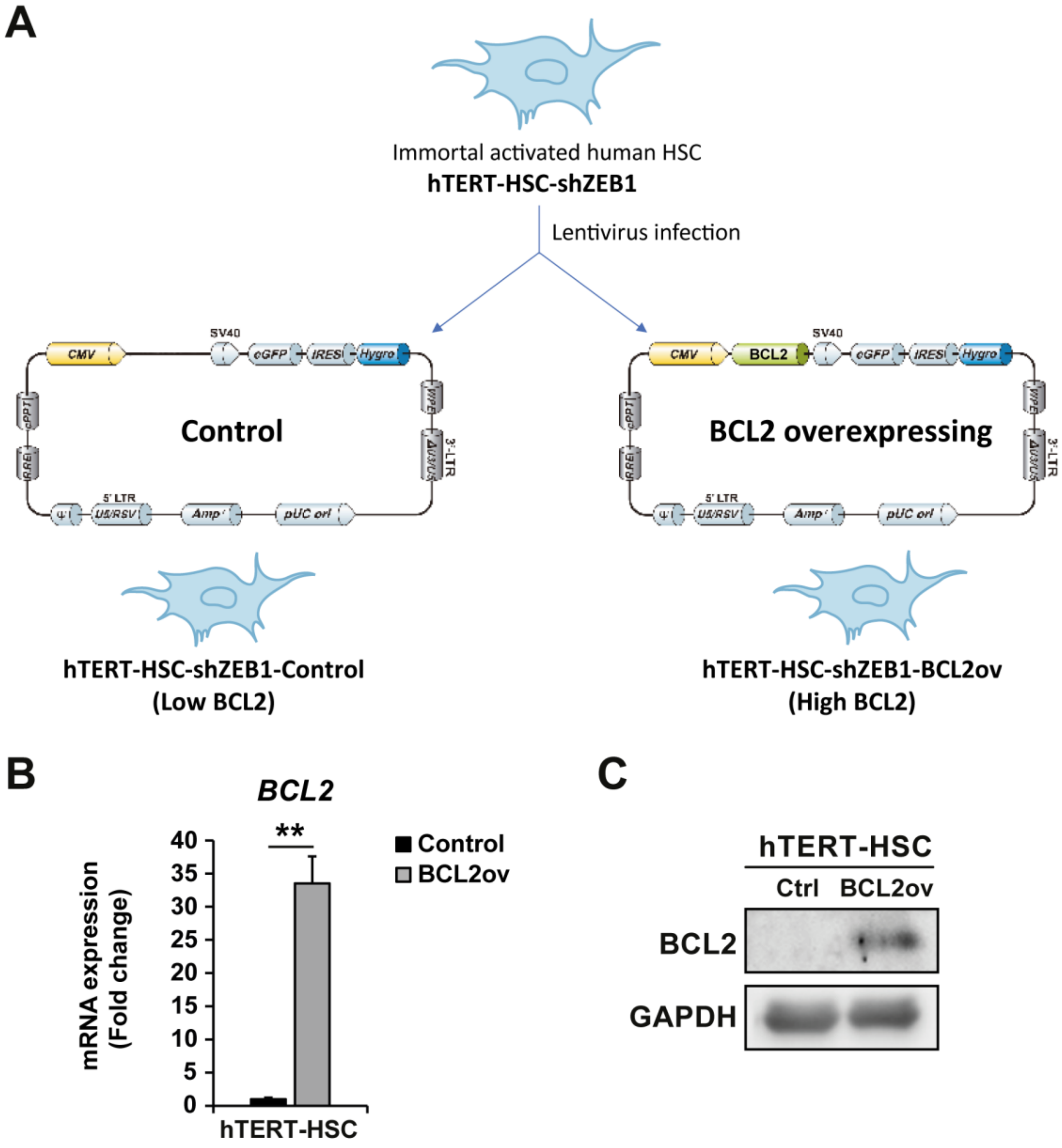


**Supplementary Figure 6. Generation of ZEB1-downregulated liver myofibroblasts with BCL2 overexpression. A.** Scheme indicating the structure of the plasmids containing the open reading frame used to produce the lentiviruses that were used to overexpress BCL2 (BCL2ov). **B-C.** Changes in BCL2 mRNA and protein expression in hTERT-HSC shRNA-ZEB1 control and BCL2ov cells, determined by RT-QPCR and Western blot. Values are expressed as means ± SEM from 5 independent cultures. **, $p<0.01$

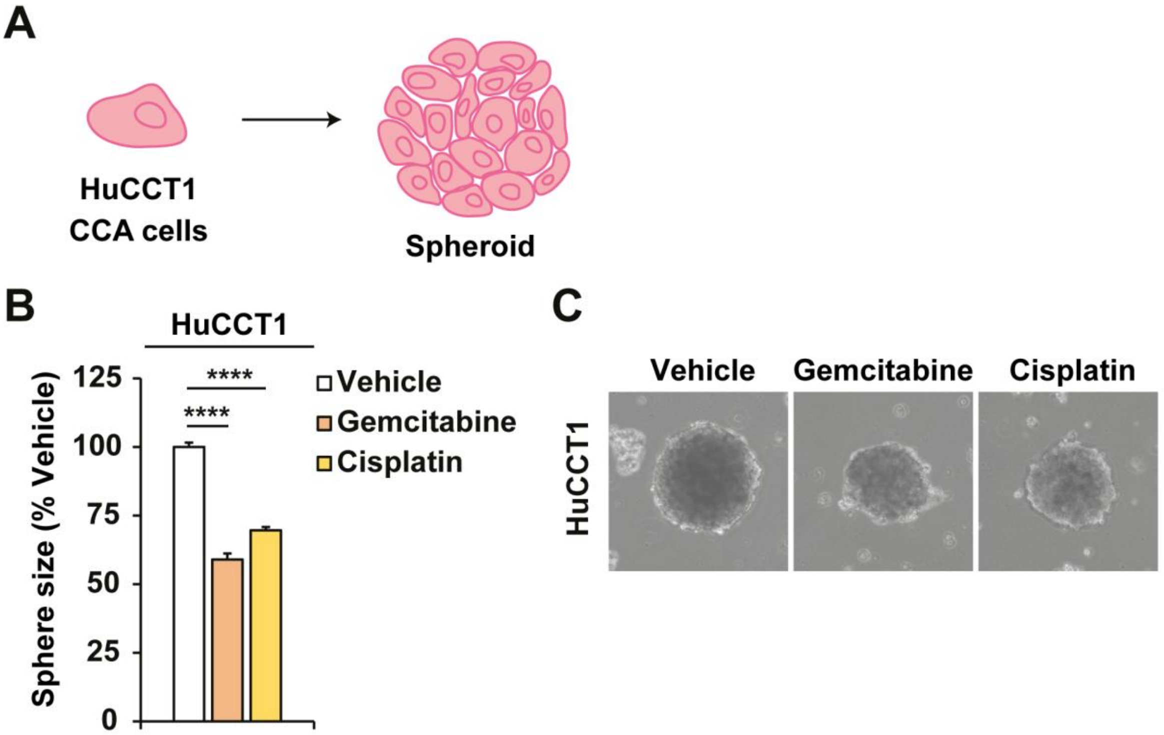


**Supplementary Figure 7. Sensitivity of tumour spheroids to chemotherapeutic drugs. A.** Scheme depicting the spheroids used in the experiments. **B.** Size of HuCCT1 spheroids treated with gemcitabine (0.1 μM) or cisplatin (25 μM). **C.** Representative images of the spheres quantified in B. Results are expressed as means ± SEM from 5 independent cultures. ****, $p < 0.0001$ as compared with vehicle.
